# UDP-6-glucose dehydrogenase in hormonally responsive breast cancers

**DOI:** 10.1101/2024.03.20.585919

**Authors:** Meghan J. Price, Annee D. Nguyen, Corinne Haines, César D. Baëta, Jovita Byemerwa, Debarati Murkajee, Sandeep Artham, Vardhman Kumar, Catherine Lavau, Suzanne Wardell, Shyni Varghese, C. Rory Goodwin

**Affiliations:** Department of Neurosurgery, Duke University Medical Center, University School of Medicine, Durham, NC, USA; Department of Medicine, John Hopkins Hospital, 1800 Orleans St, Baltimore, MD 21287, USA; Department of Molecular Genetics, Ohio State University, 1060 Carmack Road, Columbus, OH 43210, USA; Center for Population Health Sciences, Stanford University, 1701 Page Mill Road, Palo Alto, CA 94304, USA; Department of Pharmacology and Cancer Biology, Duke University Medical Center, University School of Medicine, Durham, NC, USA; Department of Biomedical Engineering, Duke University Medical Center, Durham, NC, USA; Department of Orthopedic Surgery, Duke University Medical Center, Durham, NC, USA

**Keywords:** UDP-6-glucose dehydrogenase, UGDH, breast cancer, estrogen, cancer, migration, survival, extravasation

## Abstract

Survival for metastatic breast cancer is low and thus, continued efforts to treat and prevent metastatic progression are critical. Estrogen is shown to promote aggressive phenotypes in multiple cancer models irrespective of estrogen receptor (ER) status. Similarly, UDP-Glucose 6-dehydrogenase (UGDH) a ubiquitously expressed enzyme involved in extracellular matrix precursors, as well as hormone processing increases migratory and invasive properties in cancer models. While the role of UGDH in cellular migration is defined, how it intersects with and impacts hormone signaling pathways associated with tumor progression in metastatic breast cancer has not been explored. Here we demonstrate that UGDH knockdown blunts estrogen-induced tumorigenic phenotypes (migration and colony formation) in ER+ and ER- breast cancer *in vitro*. Knockdown of UGDH also inhibits extravasation of ER- breast cancer *ex vivo*, primary tumor growth and animal survival *in vivo* in both ER+ and ER- breast cancer. We also use single cell RNA-sequencing to demonstrate that our findings translate to a human breast cancer clinical specimen. Our findings support the role of estrogen and UGDH in breast cancer progression provide a foundation for future studies to evaluate the role of UGDH in therapeutic resistance to improve outcomes and survival for breast cancer patients.

## Introduction

Accounting for 12% of new cancer diagnoses annually, breast cancer (BC) is the most frequently diagnosed cancer globally. In the United States, BC is responsible for the most cancer-related deaths after lung cancer^1^ with the leading cause of death being sequelae of metastatic disease. Survival for metastatic BC is low with a five-year survival rate of 27% for patients with estrogen receptor positive (ER+)^2,3^ and 10.81% for estrogen receptor negative (ER-) disease^4^; thus, continued efforts to treat and prevent the metastatic spread of both ER+ and ER- BC are critical. Estrogen can promote aggressive phenotypes in ER+ subtypes and increase tumorigenic phenotypes in ER- BC^5–9^ via the canonical estrogen receptor signaling pathway in ER+ tumors and alternative through growth factor signaling pathways, irrespective of ER status.

UDP-Glucose 6-dehydrogenase (UGDH) is a ubiquitously expressed enzyme critical to the formation/integration of UDP-glucuronic acid into extracellular matrix (ECM) precursors^10–12^, hormones, sugars, and xenobiotic metabolism^13, 14–17^. Dysregulation of UGDH can increase the migratory and invasive properties of ovarian cancer^16^ and triple negative BC (TNBC) ^14^ both *in vivo* and *in vitro* and is associated with stabilization of epithelial-mesenchymal transition (EMT)-associated transcription factors in lung cancer^17^. Mechanistically, the tumorigenic properties of UGDH are linked to its role in producing UDP-α-D-glucuronic acid, the precursor for glycosaminoglycans (GAGs) and proteoglycans (PGs) of the ECM and hyaluronic acids (HA)^10–12^, which are implicated in tumor progression^18,19^. While roles for UGDH in cellular migration have been defined, how it intersects with and impacts hormone signaling pathways and the pathobiology of different subtypes of metastatic BC tumors has not been explored.

Thus, we sought to assess the role of UGDH on estrogen-induced tumorigenic phenotypes in ER+ and ER- BC *in vitro* and *in vivo*. More specifically, we assess whether genetic knockdown of UGDH inhibits estrogen-stimulated migratory and invasive phenotypes *in vitro* and tumor growth and animal survival *in vivo* in both ER+ and ER- BC. We also utilize single cell RNA-sequencing to determine whether our findings translate to a human breast cancer patient using clinical specimens and patient data bases to evaluate if UGDH expression is associated with metastatic breast cancer progression.

## Results

### UGDH expression is necessary for estrogen-induced, invasive tumorigenic phenotypes in ER+ breast cancer cell lines

ER+ and ER- BC cell lines were assessed for UGDH expression, and MCF7:WS8 and T47D (ER+) or MDA-MB-231 and MDA-MB-468 (ER-) cells were used for further analysis (**Fig. 1A, Supp. Fig. 1A-I; Supp. Fig. 2A-F**). ER+ (MCF7:WS8 and T47D) cell lines stably expressing a non-targeting control shRNA (NT) and two UGDH knockdown (KD) shRNAs (U1 and U2) were generated to assess the effects of decreased UGDH expression in ER+ BC cells *in vitro* and *in vivo* **(Figs. 1B, Supp. Fig. 1A, D**). In the ER+ MCF7:WS8 line, UGDH shRNA1 (U1) effectively decreased UGDH protein expression by 90% in the presence and absence of 17β-estradiol (E2) **(Fig. 1B).** E2 increased cell migration, cell proliferation, colony formation, and cell cycle progression phenotypes in both the ER+ MCF7:WS8 and T47D cell lines *in vitro*. Wound healing and transwell migration assays showed UGDH KD significantly decreased E2-stimulated migratory phenotypes in both ER+ cell lines *in vitro* (p<0.001, **Fig. 1C**; p<0.0001, **Fig. 1D**; p<0.001, p<0.0001, **Supp. Fig. 1B, C**, E**, & F;** p<0.01). Additionally, UGDH KD altered colony formation of MCF7:WS8 and T47D *in vitro* in the presence of E2 (p<0.01, p<0.0001, **Fig. 1E, Supplemental Figure 1i, p <0.01)**. However, UGDH KD did not significantly impact E2-induced cell proliferation **(Fig. 1F)** and cell cycle progression (data not shown). UGDH knockdown, as well as decreased estrogen-mediated wound healing and transwell migration were validated via a second shRNA in both ER+ MCF7:WS8 and T47D cell lines *in vitro* (p<0.05, p<0.01, p<0.001, p<0.0001; **Supp. Fig. 1A-F**). Given the effects of UGDH knockdown on ER+ BC cells *in vitro*, we assessed whether UGDH knockdown alters primary tumor growth *in vivo.* In a mammary fat pad injection model in nude mice, knocking down UGDH significantly decreased the growth rate and reduced the final tumor size by 53% in the ER+ MCF7:WS8 cell tumors (p=0.026, **Fig. 1G; Supp. Fig. 1G)**. Of note, UGDH knockdown impacted only certain E2-induced phenotypes (migration and colony formation) but not proliferation and ERα expression **(Supp. Fig. 1H-I)**, suggesting that UGDH may affect non-canonical E2-mediated responses and/or that its effects may not depend on nuclear ERα. Therefore, we hypothesized that UGDH knockdown may impact E2-stimulated tumorigenic phenotypes in BC cell lines that do not express ERα.

**Figure 1.**
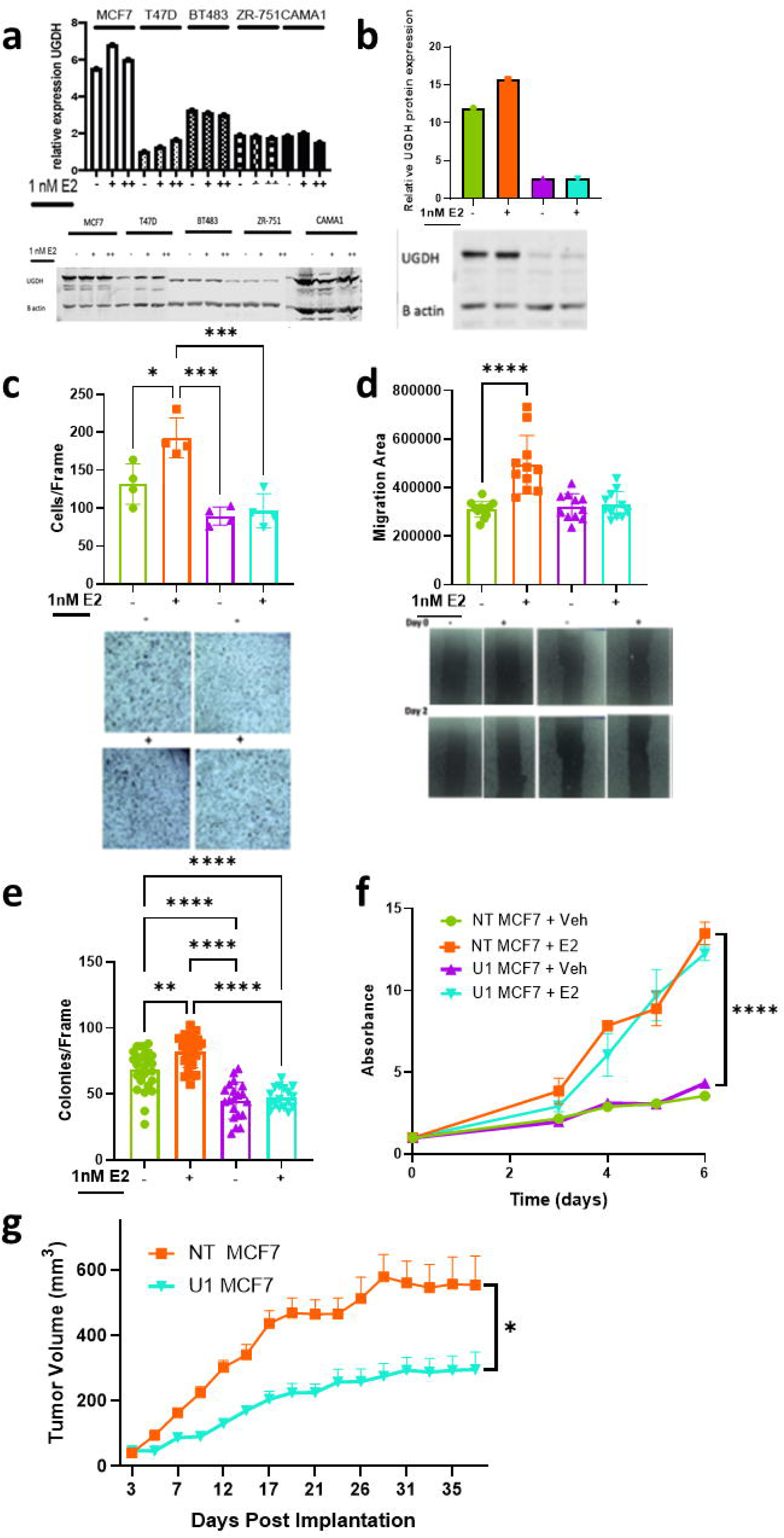
In ER+ BC cell line MCF7:WS8, UGDH knockdown significantly abrogates migratory phenotype and tumor growth in the presence (+) and absence (-) of estrogen (E2) (NT = non-targeting shRNA control; U1 = UGDH shRNA KD1. (A) Relative UGDH protein expression of ER+ breast cancer cell lines (MCF7:WS8, T47D, BT483, ZR-751, CAMA1) when treated with 1nM E2 for 24hr (+) and 48hr (++). (B) UGDH protein expression in control and UGDH knockdown cell lines, with or without E2 (NT+veh = green; NT+E2 = orange; U1+veh = purple; U1+E2 = teal). (C) Number of MCF7:WS8 cells migrating in transwell assay over 16 hours, with or without E2 stimulation, to 10% CFS (NT+veh = green; NT+E2 = orange; U1+veh = purple; U1+E2 = teal). Images at 10x magnification. (D) Differences in scratch area over 48 hours, with or without E2 stimulation on MCF7:WS8 (NT+veh = green; NT+E2 = orange; U1+veh = purple; U1+E2 = teal). Images at 10x magnification. (E) Number of MCF7:WS8 colonies formed on 1% agar after 21 days, with or without E2 stimulation. (F) Proliferation curves of MCF7:WS8 from Day 0 to Day 6, with or without E2 stimulation. (G) Tumor growth progression of subcutaneous implantation of MCF7:WS8 in mammary fat pad of Nu/J mice implanted with E2 pellets. Statistical analyses were one-way ANOVA for migration assays and colony assays; two-way ANOVA for proliferation assay and tumor growth curve, with Tukey’s post-hoc correction.*=p<0.05; **=p<0.01; ***=p<0.001; ****=p<0.0001

### UGDH knockdown abrogates estrogen-induced cancer cell migration and tumor growth independent of ER-α

The MDA-MB-231 cell line is a well validated model of TNBC which does not express ERα. MDA-MB-231 cell lines stably expressing a non-targeting control shRNA (NT) and UGDH knockdown (KD) shRNAs (U1 and U2) were generated to assess the effects of decreased UGDH expression in ERα- negative BC cells *in vitro* and *in vivo.* UGDH- shRNAs decreased UGDH protein expression by 78% and 74% in the absence and presence of E2, respectively **(Fig. 2A; Supp. Fig. 2A).** Surprisingly, migration and colony formation were significantly increased in the ERα negative cell lines (p<0.05), and as observed in ER+ cells lines, these activities were ablated upon UGDH knockdown (p<0.0001, **Fig. 2B-C**; p<0.01, **Fig. 2D**; p<0.05; p<0.001, p<0.0001, **Supp. Fig. 2B-C)**. No effects on proliferation were noted under any of the conditions tested **(Fig. 2E).** UGDH knockdown, as well as decreased estrogen- mediated wound healing and transwell migration were validated via a second shRNA in ER- MDA-MB- 231 (p<0.05, p<0.01, p<0.001, p<0.0001; **Supp. Fig. 2A-C**) and MDA-MB-468 cell lines (p<0.05, p<0.01, p<0.001, p<0.0001; **Supp. Fig. 2D-F**). Given the effects of UGDH knockdown were seen in both ER+ and ER- cell lines, we investigated the effects of UGDH on other signaling pathways parallel to estrogen receptor signaling, such as the membrane bound G-protein coupled estrogen receptor (GPER30/GPER). Our investigation into the GPR30/GPER pathway showed a decrease in GPR30 expression in estrogen treated UGDH knockdown lines that correlated to decreased migratory phenotypes via wound healing and transwell migration in MCF7 and MDA-MB-231 (p<0.05, p<0.01, p<0.001, p<0.0001; **Supp. Fig. 3A-F**).

**Figure 2.**
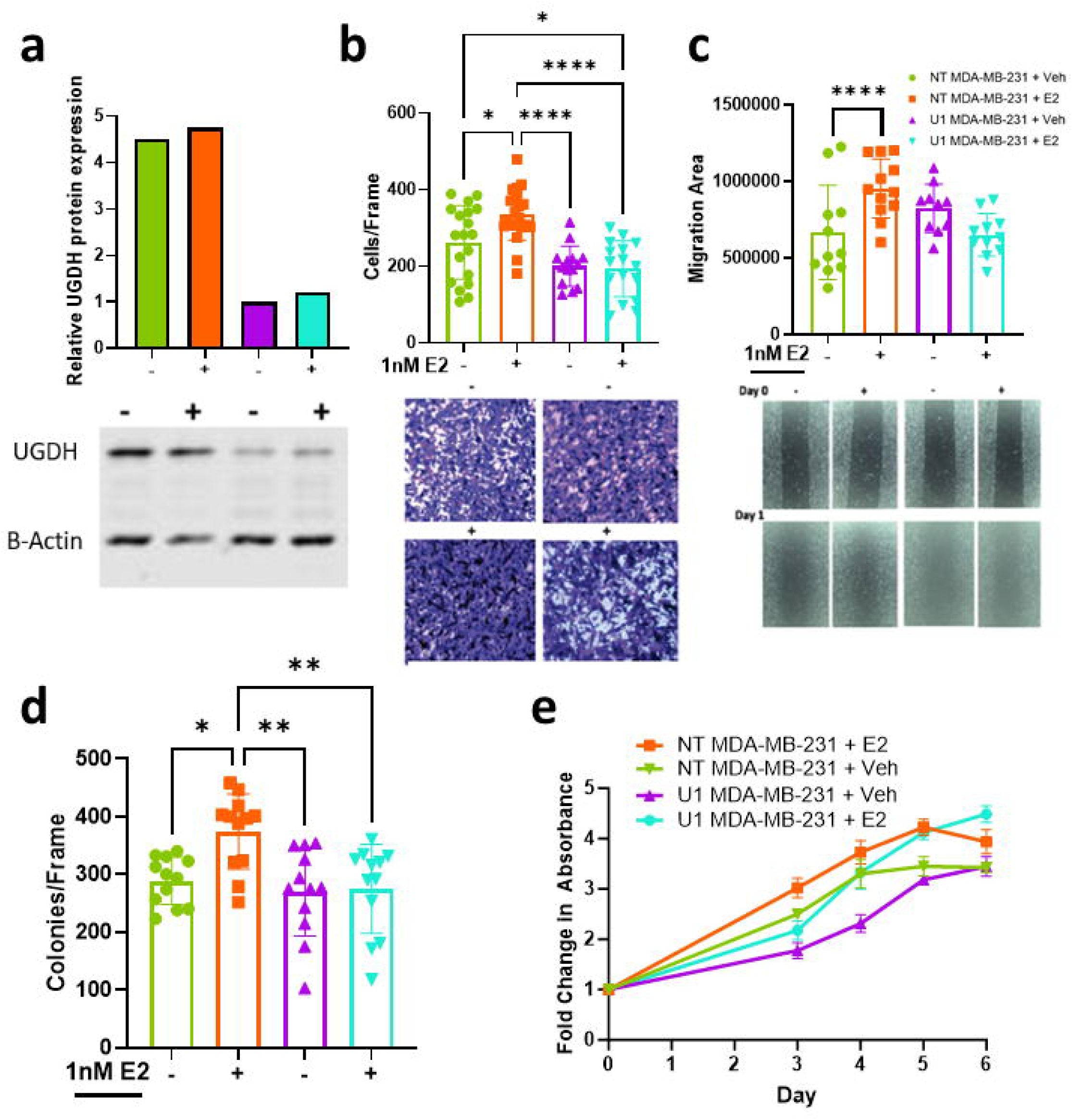
In ER- BC cell line MDA-MB-231, UGDH knockdown significantly abrogates migratory phenotype and tumor growth in the presence (+) and absence (-) of estrogen (E2) (NT = non-targeting shRNA control; U1 = UGDH shRNA KD1). (A) UGDH protein expression in NT control (green and orange) and UGDH knockdown (purple and teal) cell lines, with or without E2. (NT+veh = green; NT+E2 = orange; U1+veh = purple; U1+E2 = teal) **(B)** Number of MDA-MB-231 cells migrating in transwell assay over 16 hours, with or without E2 stimulation, to 10% CFS (NT+veh = green; NT+E2 = orange; U1+veh = purple; U1+E2 = teal). Images at 10x magnification. **(C)** Differences in scratch area over 48 hours, with or without E2 stimulation on MDA-MB-231 (NT+veh = green; NT+E2 = orange; U1+veh = purple; U1+E2 = teal). Images at 10x magnification. **(D)** Number of MDA-MB-231 colonies formed on 1% agar after 21 days, with or without E2 stimulation. **(E)** Proliferation curves of MDA-MB- 231 from Day 0 to Day 6, with or without E2 stimulation. *=p<0.05; **=p<0.01; ***=p<0.001; ****=p<0.0001. Statistical analyses were one-way ANOVA for migration assays and colony assays; two-way ANOVA for proliferation assay and tumor growth curve, with Tukey’s post-hoc correction.

### UGDH knockdown impairs extravasation of ER- breast cancer ex vivo, regardless of E2 stimulation

UGDH plays a critical role in the tumor microenvironment and prior studies demonstrate that it may be implicated in metastasis. Given the estrogen-mediated migration and invasion phenotype in vitro, we sought to examine the effects of UGDH knockdown on the initial steps of BC metastasis, using MDA- MB-231 NT shRNA or UGDH shRNA cell lines perfused through an *ex vivo* microfluidic chip with endothelial vasculature to model human microcirculation and extravasation during the metastatic cascade **(Fig. 3A).** UGDH knockdown significantly reduced percentage of extravasating MDA-MB-231 cells, regardless of estrogen stimulation (∼16-fold; p = 0.0284) **(Fig. 3B)**.

**Figure 3.**
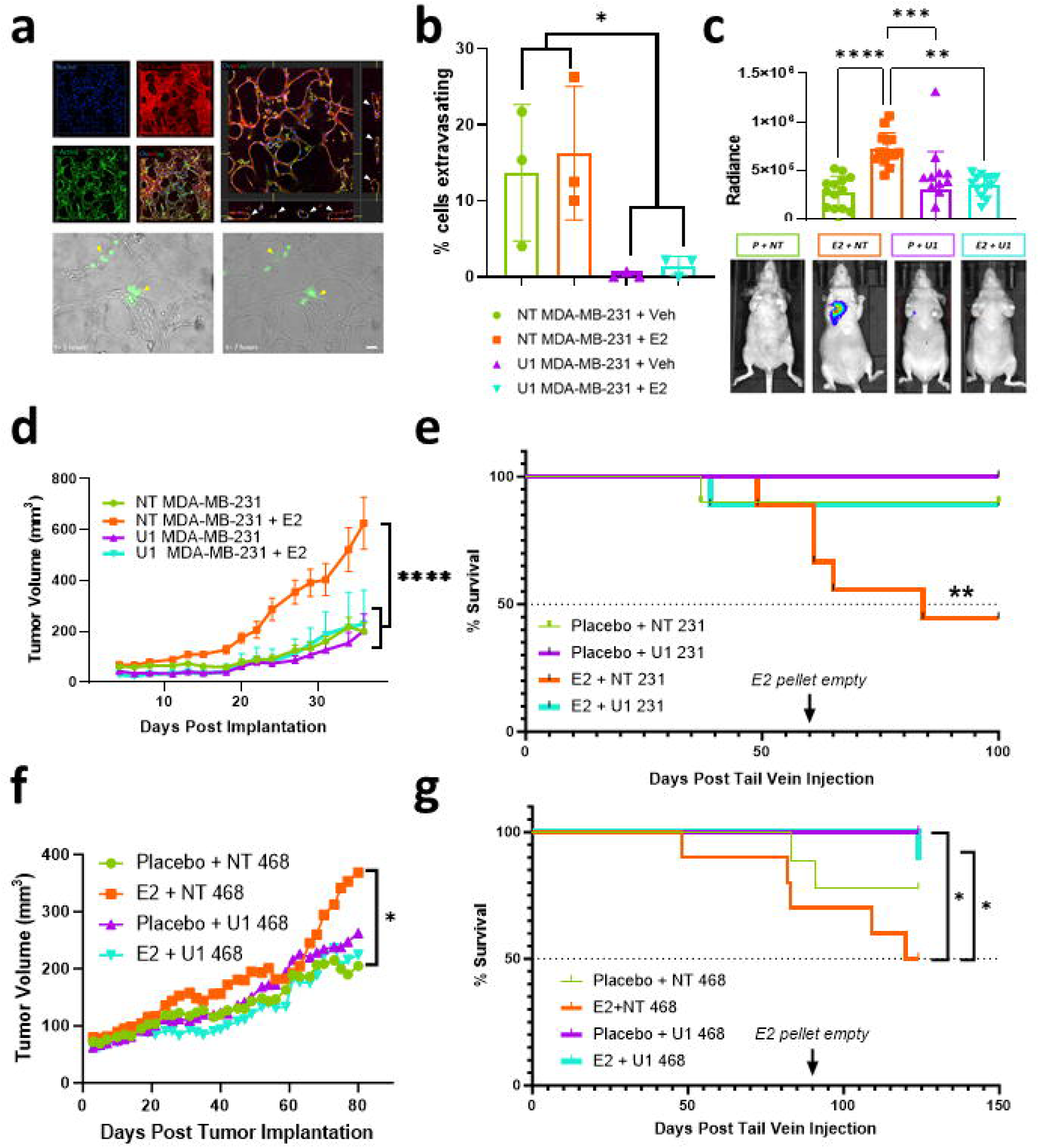
UGDH is associated with estrogen-mediated metastatic breast cancer progression in ER- breast cancer and with decreased survival in vivo. UGDH knockdown impairs metastasis by preventing extravasation of MDA-MB-231 cells ex vivo, even in the presence of estrogen (1nM). (A) Characterization of the tissue-engineered vasculature network using staining for VE-Cadherin, F-actin, and nuclei. Orthogonal views of z-stack confocal images of the immunostained vasculature showing vascular lumens (white arrows). MDA-MB-231 cells (green) extravasating out of the endothelial vasculature network over time. Extravasating cellSStatiss indicated with yellow arrows. Images scale: 50 um. (B) Percent (%) of cells extravasating ex vivo with or without E2 stimulation in MDA-MB-231 cells. (C) Representative bioluminescent images of mice treated with estrogen or placebo pellets and implanted with NT shRNA or U1 shRNA MDA-MB-231 luciferase (Luc) labeled cells and radiance values of tumor burden at Week 7 (peak signal). (D) Tumor growth progression of subcutaneous implantation of MDA-MB-231 in mammary fat pad of Nu/J mice. (E) Survival curve of metastasis experiment with mice treated with estrogen or placebo pellets and implanted with NT shRNA (NT) or U1 shRNA MDA-MB- 231 (U1) luciferase (Luc) labeled cells. (F) Primary mammary fat pad xenograft tumor growth curve of Nu/J animals implanted with NT shRNA or U1 shRNA MDA-MB-468 cells treated with placebo or E2 pellets. () Survival curve of metastasis experiment with mice treated with estrogen or placebo pellets and implanted with NT shRNA (NT) or U1 shRNA MDA-MB-468 luciferase (luc) labeled cells. Statistical analyses were student T-test with Bonferroni post-hoc correction for ex vivo analysis; one-way ANOVA for radiance analysis & tumor growth curve, with Tukey’s post-hoc correction; and Gehan-Wilcoxon test with Sidak’s post-hoc correction for survival curves. *=p<0.05; **=p<0.01; ***=p<0.001; ****=p<0.0001.

### UGDH knockdown impairs tumor growth and metastatic progression of ER-negative breast cancer cells

### in vivo

To examine the effect(s) of UGDH knockdown on metastatic tumor progression *in vivo,* Nu/J mice implanted with either placebo (P) or estrogen pellets (E2) were injected via tail vein with either Control (NT) or UGDH shRNA1 (U1) MDA-MB-231 cells. E2+NT mice showed the greatest metastatic tumor burden compared to P+NT (p<0.0001); P+U1 (p=0.0001); and E2+U1 (p=0.0011) (**Fig. 3C)**. To investigate the effect of E2 stimulation and UGDH KD on primary ER- tumor growth, we injected control shRNA or UGDH knockdown shRNA MDA-MB-231 cells into the mammary fat pad of mice. E2-stimulated NT shRNA MDA-MB-231 tumors demonstrated the greatest growth rate and size (307% of unstimulated tumors at Day 36, p < 0.0001)). Interestingly, UGDH knockdown completely abrogated the tumor growth- promoting effects of E2 **(Fig. 3D).** Given that UGDH knockdown blunted the E2-induced primary tumor growth response *in vivo* and significantly reduced extravasation in the microfluidic *ex vivo* model, we hypothesized that UGDH knockdown would blunt metastatic progression and increase animal survival. All U1-injected mice (P+U1 and E2+U1) and P+NT mice survived significantly longer (MS undefined, p=0.0101) than E2+NT mice (MS = 84 days) (**Fig. 3E**). To confirm that the estrogen-mediated tumor progression in ER- model *in vivo* was not cell-line specific, MDA-MB-468 cells were injected via mammary fat pad or tail vein with either Control (NT) or UGDH shRNAs to assess primary ER- tumor growth, metastatic progression and animal survival. Again, E2-stimulation was associated with increased primary tumor growth and resulted in worse overall survival in ER- BC *in vivo*, and these effects were abrogated by UGDH KD (p<0.05; **Fig. 3F-G, Supplemental Figure 4**).

### UGDH is involved in metastatic breast cancer progression in a patient

Approximately 70% of advanced breast cancer patients develop bone metastasis with the spine being the most common site^20,21^ and recent studies indicate that breast cancer has a metastatic tropism to the spinal vertebral body^22^. To determine whether our findings could translate to a human breast cancer patient, single-cell RNA-sequencing of clinical specimens obtained from a radiologically normal and tumor-containing vertebral body (**Fig. 4A-D**) from the same patient was performed, using our established protocol^23^, to determine whether UGDH expression was associated with metastatic breast cancer progression. Biopsied specimens obtained from the corresponding vertebral bodies were verified by immunohistochemistry (**Fig. 4E & 4F**), confirmed to be breast in origin (**Fig. 4G-I**) and processed for single cell sequencing. UMAP analyses showed UGDH was upregulated in tumor, compared to normal vertebral tissue, in cancer epithelial cells (**Fig. 4J-K**). After observing that UGDH was upregulated in metastatic breast cancer human clinical specimens, the association between UGDH expression level and recurrence free survival (RFS) in patients with high grade ER+ and ER- BC was assessed. Higher UGDH protein expression correlated with worse prognoses for high grade ER+ BC and ER- BC (p = 0.043; 0.0038) **(Fig. 4L-M).**

**Figure 4.**
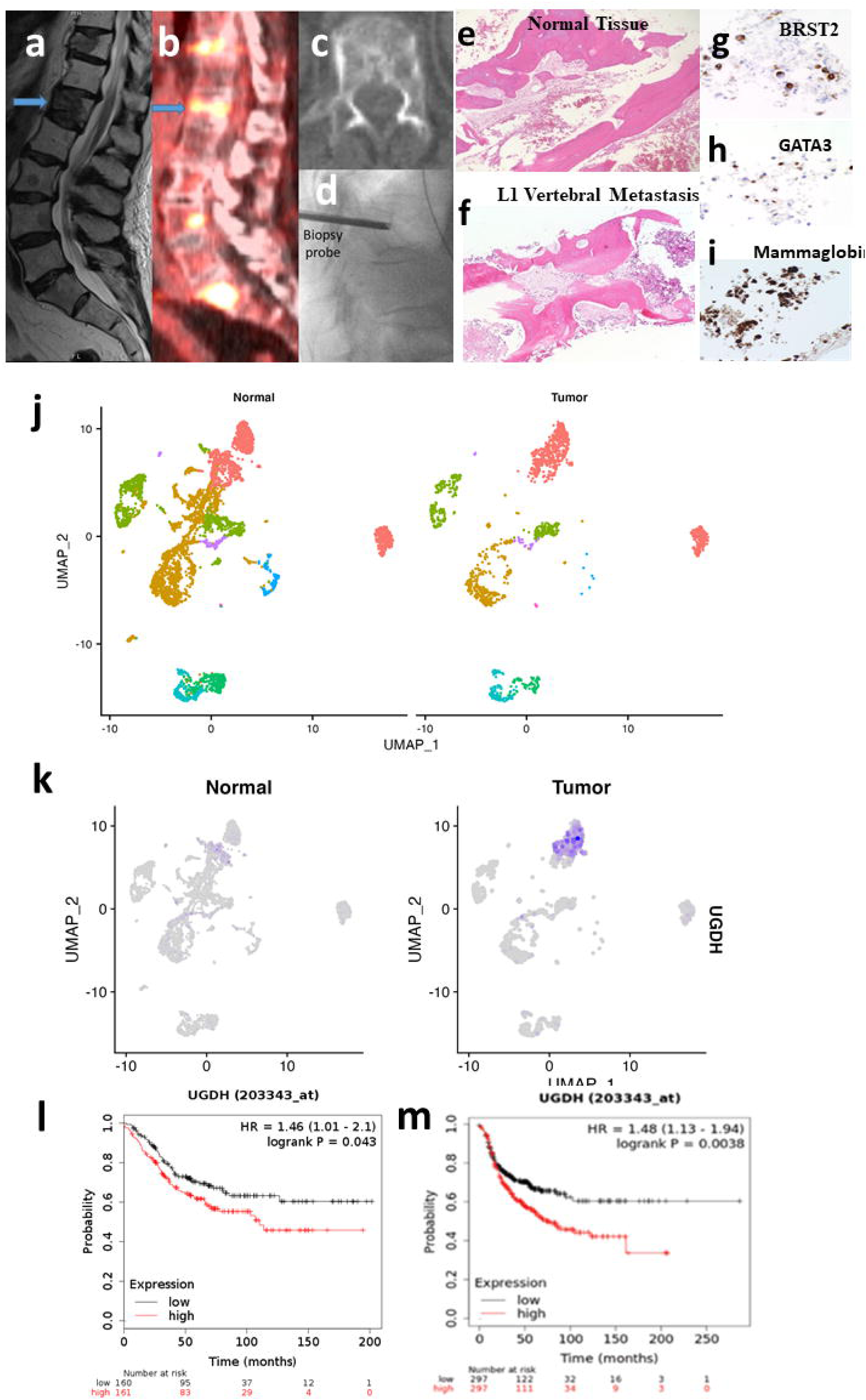
UGDH is associated with metastatic breast cancer progression in patients and with lower progression free survival. (A) Pre-operative sagittal MRI, **(B)** sagittal PET scan, **(C)** axial CT, and **(D)** intraoperative X-ray of L1 breast cancer spine metastasis (blue arrow pointing to metastatic mass in the vertebral body). **(E)** Histology showing H&E of normal vertebral tissue and **(F)** H&E of L1 vertebral metastasis. **(G)** Immunohistochemistry of BRST2, **(H)** GATA3, and **(I)** mammaglobin to demonstrate that metastatic tumor is breast in origin. **(J)** UMAP depicting 26 unique cell subset clusters for normal and tumorous tissue in a metastatic breast cancer patient. **(K)** UMAP depicting UGDH expression in normal and tumorous tissue, concentrating in epithelial cells with high copy number variations. **(L)** UGDH expression in ER+ BC correlated to lower recurrence free survival. **(M)** UGDH expression in ER- BC correlated to lower recurrence free survival.

## Discussion

Our study establishes the involvement of UGDH in estrogen-induced breast cancer progression and also provides further evidence of the role of estrogen in the development of metastatic progression, irrespective of ER status. Specifically, we demonstrate that estrogen induces migratory and invasive phenotypes in both ER+ and ER- BC *in vitro*, which are abrogated by UGDH knockdown. Additionally, UGDH knockdown significantly reduces estrogen-dependent tumor growth in ER+ and ER- primary BC orthotopic models *in vivo*. *Ex vivo* microfluidics model demonstrated that UGDH can regulate extravasation in the presence and absence of estrogen. Also, UGDH knockdown blunted the metastatic formation and progression of ER- BC lines stimulated by estrogen in tail vein models of metastasis *in vivo*. Furthermore, in a human breast cancer patient, UGDH expression was upregulated in metastatic breast cancer tissue compared to normal vertebral tissue in a patient, specifically in cancer epithelial cells. In the context of our *in vitro*, *in vivo*, and clinical explorations, UGDH appears to be involved in metastatic breast cancer progression and mitigating UGDH expression can abrogate migratory phenotypes that may contribute to hormonally responsive metastatic breast cancer.

To our knowledge, this is the first study demonstrating that UGDH expression can mediate the tumorigenic effects of hormone stimulation via estrogen in BC regardless of ER status. Estrogen is known to increase the risk of BC development^24^ via activation of estrogen receptor alpha (ERα) in ER+ BC with subsequent upregulation of genes associated with tumor cell invasion, growth and survival^25^. While not widely considered to be estrogen responsive, previous literature^26^ suggests that ER- cell lines can become more aggressive in the presence of estrogen, which is consistent with our findings.

Mechanistically, crosstalk between estrogen stimulation, growth factor signaling pathways involving receptor tyrosine kinases such as IGF-1R and EGFR, and cell surface receptors including G protein- coupled receptors (GPCRs) and their downstream mechanisms through the PI3K/AKT and MEK/ERK pathway have been implicated in estrogen effects that are ER independent^27–31,26,32,33^. IGF-1R signaling is also involved in EMT-mediated metastatic tumor progression and IGF-1R overexpression is shown to promote migratory phenotypes of TNBCs via the focal adhesion kinase signaling pathway^32,33^. Of the noted ER-independent pathways, UGDH has been shown to interact with and modulate the PI3K/AKT and the MEK/ERK pathways downstream of growth factor receptors in other cancers through which cancer cell motility, invasion, migration, and survival can occur ^19, 33, 34^.

Congruent with our findings, previous studies also demonstrated that UGDH knockdown abrogates tumor growth, invasion, and colony formation but does not impact proliferation in TNBC cell lines^14,34^, glioblastoma (GBM)^15^, and lung adenocarcinoma^17^. Conversely, UGDH knockdown can mitigate both tumor cell proliferation and migration^35^ in colorectal carcinoma and in ovarian cancer^16^. Although our findings align with the previously proposed “go-or-grow” hypothesis that cancer cell motility and proliferation can be mutually exclusive, ^36,37^ it is important to note that this dichotomy is not ubiquitous across all cancer types. Although further investigations are needed to determine the precise mechanism through which UGDH may modulate the estrogen-induced effect on migration and tumor growth, targeting UGDH can abrogate estrogen-induced tumor phenotypes, regardless of ER status.

Aside from affecting canonical cancer signaling pathways, UGDH may also promote estrogen-mediated tumor progression by influencing cancer cell motility and/or the associated tumor microenvironment, as specifically seen in our exploration of the tumor microenvironment of a patient with metastatic breast cancer in which UGDH was highly expressed in tumor epithelial cells and was absent from normal vertebral tissue. Dysregulation in the UGDH-driven GAG/PG production pathway has been described as a possible explanation for UGDH knockdowns impairment of cancer cell motility^14^. Taking our results in the context of recent GBM studies, UGDH may aid in generating extracellular matrix precursors that influence cancer cell migration, EMT processes, and macrophage-related transcriptional, and epigenetic regulatory pathways to modulate estrogen-induced tumorigenic phenotypes^38^. Thus, further investigation into the role of UGDH on EMT, ECM production, and hormonal metabolism in the breast cancer tumor microenvironment are also warranted to fully elucidate its role in breast cancer progression.

## Conclusion

Our study is the first to evaluate the role of UGDH in hormonally responsive breast cancers (BC). We have established that UGDH knockdown effectively mitigates estrogen stimulated phenotypes in BC *in vitro* and *in vivo* irrespective of estrogen receptor status. These studies support the role of estrogen and UGDH in breast cancer progression provide a foundation for future studies to evaluate the role of UGDH in therapeutic resistance to improve outcomes and survival for breast cancer patients.

## ACKNOWLEDGMENTS

We thank Ching-Yi Chang and Donald McDonnell for providing guidance and expertise in interpretation of results for all experiments.

## Methods

### Reagents and Cell Culture

All reagents were purchased from Sigma-Aldrich unless otherwise specified. Puromycin was diluted to a concentration of 1 ug/mL in cell culture medium as a working concentration. MCF7:WS8, T47D, and MDA-MB-231 (American Type Culture Collection, ATCC, Manassas, Virginia) were grown in Dulbecco’s Modified Eagle Medium (DMEM), Roswell Park Memorial Institute (RPMI), and Minimum Essential Media (MEM) respectively, all supplemented with 10% fetal bovine serum (FBS), non- essential amino acids (1%), sodium pyruvate (1%), and puromycin. 17-β-estradiol (E2) (Cat# E-060, Sigma Aldrich) was used in all assays at a physiologic concentration of 1 nM/mL. To assess the effect of hormonal stimulation, twice charcoal stripped media at either 10% or 0.5% was used for all assays evaluating the role of E2. G1 (GPER/GPR30 agonist) from Tocris Bioscience was used at 0.1 uM/mL, and G15 (GPER/GPR30 antagonist) from Tocris Bioscience was used at 1 uM/mL. To appropriately assess the effect of hormonal stimulation, twice charcoal stripped media (McDonnell lab) at either 10% or 0.5% was used for all assays evaluating the role of E2 and G1 stimulation.

### Lentiviral Transduction

UGDH shRNA lentiviral particles were purchased from Dharmacon (Buckinghamshire, UK) as a set of 3 SMARTvector Inducible Human UGDH shRNA (Cat # V3SH7675-01EG7358). Control (non-targeting) shRNA clone (Cat # VSC6570), UGDH shRNA1 (Cat # V2LHS_171838), and UGDH shRNA2 (Cat # V2LHS_218865) were used. MCF7, T47D, and MDA-MB-231 cells were transduced with virus in polybrene media (8 ug/mL) for 48 hours prior to puromycin selection (1 ug/mL) as previously described (Xia et al, 2016).

### Quantitative real-time PCR

Total RNA was extracted via QIAquick PCR Purification Kit (Qiagen, Mansfield, MA). After reverse transcription using iScript cDNA Synthesis Kit (Bio-Rad, Hercules, CA) and Oligo(dT) primer, quantitative real-time PCR (qRT-PCR) was performed using SYBR Green PCR Mix (BioRad, Hercules, CA) and IQ5 detection system (Bio-Rad, Hercules, CA). Relative gene expression was normalized to 36B4 or ACTB gene expression.

### Immunoblot and Immunocytochemistry

Total cellular protein was extracted with radioimmunoprecipitation assay (RIPA) buffer containing protease, phosphatase inhibitors and sodium orthovanadate. SDS-PAGE was performed with 50 ug total proteins using 10% gradient acrylamide gels (LI-COR Biosciences, Lincoln, NE). Western blot analysis was performed using Quantitative Western Blot System, with secondary antibodies labeled by IRDye infrared dyes (LI-COR Biosciences, Lincoln, NE). Antibodies included: anti-UGDH (ab15505, Abcam, Cambridge, MA), and anti-GPER/GPR30 (AF5534, R&D Systems, Minneapolis, MN), and anti- β-actin (8H10D10, Cell Signaling, Danvers, MA).

### Proliferation Assay

2,000 cells per well were plated in 96 well plate and starved in 10% charcoal stripped (CS) FBS- supplemented media for 48 hours and then stimulated with either 1 nM E2 or continued 10% CS media. Baseline CellTiterGlo (Promega) reading was taken at specified time points.

### Cell Cycle Analysis

Cell cycle was analyzed by flow cytometry. Cells were plated in 10% CS medium for 48 hours followed by stimulation with either E2 or nothing for the indicated time points. Cells were then trypsinized and dissociated by pipetting, fixed with 75% ethanol at 4 C for 30 min. Cells were incubated with DNase free RNase at 37 C for 30 min followed by propidium iodide (100 ng/ml) for 1 h at 37 C. Percentage of cells at each phase (G1/G0, S, and G2/M) were analyzed using BD CellQuest Pro Software (Becton Drive Franklin Lakes, NJ).

### Wound Healing Assay

Cells were grown in 10% CS serum in 6 well plates until confluent. Three scratches per well were created using a wide-tip 10 uL pipette tip through the confluent cells. Dishes were washed with PBS and cells were grown in 0.5% CS medium with or without estrogen stimulation for 24-48 hours. Phase contrast pictures were taken at 0, 24, and 48 hours. The width of the scratch was measured and quantified using the MRI Wound Healing plugin in ImageJ.

### Transwell Migration Assay

Cells were grown in 10% CS serum in 6 well plates for 24 hours; they were then either starved with 0.5% CS (control) or treated with 1 nM E2, for 24 hours. Transwell inserts (COSTAR) were placed into twenty-four well plates and were seeded at 50,000-75,000 cells per insert in 100 uL of either 0.5% CS media or 0.5% CS media supplemented +/- E2 for 24 hours. Cells migrated towards media with 10% CFS for 12-24 hours and then were fixed with 30% formaldehyde, stained with Crystal Violet and then washed with phosphate buffered saline (PBS). Pictures of the entire transwell surface were taken, and cell counts were quantified using ImageJ.

### Ex Vivo Microfluidic Metastasis Model

The microfluidic vasculature-on-a-chip device was used to assess cancer cell extravasation *ex vivo*. The device was fabricated using soft lithography as reported ^37,38^. Briefly, SU-8 100 photoresist (Microchem) was spun on a 4 inch silicon wafer (University Wafers) and cured via light exposure through a photomask to create features with a height of 100 µm. The uncured SU-8 was washed away using SU-8 developer and the wafer was silanized using Trichloro(1H,1H,2H,2H-perfluorooctyl)silane (Sigma 448931). Sylgard 184 (Ellsworth) polydimethylsiloxane (PDMS) at 10:1 (base:crosslinker) was poured onto the wafer and allowed to cute at 60 °C for 2 hours. Post curing, the PDMS was cut around the features, inlets and outlets were punched using biopsy punches, and bonded to a cover glass using plasma treatment for 60 s.

The device consisted of three parallel microchannels with an array of trapezoidal microposts separating the three channels. Human umbilical vein endothelial cells (Lonza, C2519A) were suspended in 4 units/ml thrombin solution at a concentration of 50 million/ml, mixed 1:1 with 5 mg/ml fibrinogen solution, and perfused into the central microchannel of the device to allow gelation. After gelation, media channels were flushed with cell culture medium and the devices were cultured for 5 days to allow the endothelial cells to self-assemble into microvascular networks with perfusable lumens. The engineered tissue for immunostained for VE-Cadherin (Bio-techne, AF938) (1:100 dilution) followed by donkey anti-goat (1:200) secondary antibody to visualize the endothelial cell-cell junctions. Alexa Fluor 488 phalloidin (Thermo Fisher, A12379) and Hoechst 33342 were used to stain actin and nuclei respectively. Fluorescently labelled control (NT shRNA) and UGDH knockdown (UGDH KD1) MDA-MB-231 cell lines were suspended in medium and perfused into the vascular networks in the presence or absence of estrogen (1nM) and the microfluidic devices were cultured for up to 8 hours to allow the cells to extravasate from the vasculature into the fibrin hydrogel. Number of extravasating cells were assessed per condition using ImageJ software.

### Animal Studies

#### A. Mammary Fat Pad Models

All protocols involving animals were previously approved by the Institutional Animal Care and Use Committees. Mice used in all experiments were female Nu/J (Cat# 002019, Jackson Laboratory, Bar Harbor, ME) aged 6-8 weeks. Sample size was determined by end-point statistics (power analysis) performed prior to study initiation by Dr. James Herdon (N = 30). For MCF7:WS8 experiments mice were ovariectomized and implanted with 17B-estradiol pellets (0.36mg, 60 days, Cat# SE-121, Innovative Research of America). Mice then received mammary fat pad injections of 1.3 x 10^6^ viable MCF7:WS8 cells that were stably transfected with the non-targeting (NT) shRNA, UGDH shRNA1 or UGDH shRNA3. Tumor volumes were measured three times weekly for 5 weeks. Tumor volumes were calculated using the formula (short width * short width * long width)/ 2. After sacrificing mice, primary tumors were removed and processed for RNA and protein analysis.

The same protocol was used for the mammary fat pad experiments with MDA-MD-231 and MDA-MB-468 with the following modifications. Mice (N=40) were ovariectomized and implanted with either placebo pellets (Cat# SC-111, Innovative Research of America) (groups 1 and 3) or 17β-estradiol pellets (groups 2 and 4). Groups 1 and 2 were then injected with NT shRNA MDA-MB-231 cells into the mammary fat pad while groups 3 and 4 received UGDH shRNA1 MDA-MB-231 cells. The resulting experimental groups were the following (N=10 mice/group): (1) NT shRNA + placebo; (2) NT shRNA + estrogen; (3) UGDH KD1 + placebo; (4) UGDH KD1 + estrogen. Tumor volume measurement and tissue harvest were the same as described above.

#### B. Metastasis Models

To establish the metastatic BC models, mice (N=60) were ovariectomized and implanted with either 17β-estradiol pellets or placebo pellets. Parental cells used to generate the luciferase labeled and shRNA lines below were MDA-MB-231 and MDA-MB-468 (ATCC, Duke University). Mice then received arterial tail vein injections of 1 x 10^6^ viable cells expressing luciferase (Luc) and infrared protein (iRFP) that were stably transfected with NT shRNA or UGDH shRNA1. The resulting experimental groups were the following (N=15 mice/group): (1) placebo pellet + NT shRNA MDA-MB-231 or MDA-MB-468 (P+NT); (2) estrogen pellet + NT shRNA MDA-MB-231 or MDA-MB-468 (E2+NT); (3) placebo pellet + UGDH KD1 MDA-MB-231or MDA-MB-468 (P+U1); (4) estrogen pellet + UGDH KD1 MDA-MB-231 or MDA-MB-468 (E2+U1).

To track metastatic tumor progression, mice were imaged weekly using the IVIS Lumina III In Vivo Imaging System (PerkinElmer, USA) to quantify tumor burden as normalized radiance. Prior to imaging, mice were first injected via intraperitoneal administration with 100uL of 15 mg/mL D-Luciferin Sodium Salt (Cat# 1-360243-200, Regis Technologies, USA) reconstituted in sterile 1X Dulbecco’s PBS. Images were collected at exposure times of 1 second, 30 seconds, 60 seconds, and 180 seconds and quantified as normalized radiance.

Mice were weighed and monitored three times weekly for survival until end of the study. Mice were euthanized when body weight fell below 15% of their original Day 0 weights or when mice could no longer ambulate for food or water due to tumor-related paralysis. Upon median survival of any group, N=5 mice per group were euthanized and harvested for lung, liver, and uterus to be weighed and noted for tumor burden and estrogen effects. Mice harvested for the median survival time point or found dead due to non-tumor-related causes before week 1 of the study were censored from the final survival curve.

### Cell Line Sequencing

Gene expression analysis from cell lines were generated by mRNA sequencing using Illumina NovaSeq 6000. Briefly, 1 ug of total RNA was converted to RNA sequencing libraries using the Kapa Stranded mRNA-seq library prep kit (Manufacture, Location) and sequencing on an Illumina NovaSeq 6000 using the 50bp paired-end configuration with an average of 51.2 million read pairs per sample. Reads were aligned to GRCh38 using STAR (PMID: 23104886). Transcript abundance estimates were performed for each sample using Salmon (PMID: 28263959). Differential expression analysis was performed using DESeq2 (PMID: 25516281) with an FDR cutoff of 5%.

### Single Cell Sequencing of Patient Specimens

#### A. Patient History

All experiments for this study are performed under the IRB protocol Pro#00101198. The patient was a 71- year-old female with a history of ER+/PR+/Her2 negative metastatic breast cancer status post T10-L2 radiation to the spine 30Gy in 10 fractions completed three months prior to the surgical procedure. Patient was started on palbociclib one month later in August 2020, held for 2 weeks during August for a root canal/tooth abscess and re-initiated later that month (August 24, 2020). Patient was receiving palbociclib consistently until October 10, 2020 when a PET scan and MRI findings identified progression of disease at T11 and L1 corresponding to PET scan (Fig. 1A-C). The patient was taken to the OR for biopsy/radiofrequency ablation and kyphoplasty with cement-augmentation.

#### B. Surgical Procedure for Specimen Acquisition

After patient was deemed a surgical candidate based on clinical indications and consent obtained for the collection of tissue, the patient is placed under general anesthesia, the vertebral body (T11-L1) is identified with fluoroscopic guidance. The skin overlying the target area is prepped with chlorhexidine and sterilely draped in the usual fashion maintaining meticulous sterile technique. Local anesthesia is provided with 1% Lidocaine. A stab incision is made 1 cm lateral of the lateral border of each pedicle at the level of the target vertebral body. A trocar is introduced safely through each pedicle just inside the target vertebral body. The stylet of the trocar is removed and a working cannula is left in place. The biopsy cannula is inserted, and anterior/posterior and lateral fluoroscopic x-rays views are taken to verify accurate placement of biopsy cannula (Fig. 1D). A biopsy is taken from the representative levels, and divided in half. One half of the specimen is sent to pathology for further evaluation to confirm presence of normal or tumor specimen (Fig. 1E & F) and evaluated for its tissue of origin (Fig. 1G-I). The other half is placed in a pathology specimen cup and taken to the laboratory for further single cell sequencing processing within 30 minutes of acquisition, preferably transferred on ice to maintain cell viability. The remainder of the procedure is performed according to clinical indications and then closed in the standard fashion.

#### C. Sample Processing for Single Cell Studies

Human samples were processed within a biological safety cabinet. Tissues from tumor and normal vertebral bodies were manually dissected from the block of tissue using razor blades, surgical scissors, and forceps, to obtain pieces of tissues that are less than 2 grams total. Samples are then minced using a razor blade and placed into a gentleMACS C tube (Miltenyl Biotec, cat.no. 130-093-237) containing serum-free DMEM, 1 mg/mL Collagenase A (Millipore Sigma, cat.no. 10103586001), and 0.1 mg/mL DNAse I (Sigma-Aldrich, cat.no. D5025-150KU). Samples are homogenized on a gentleMACS Dissociator (Miltenyl Biotec, cat.no. 130-093-235) with two rounds of the “h_impTumor_02.01” program and then placed into a shaking incubator for 30 minutes at 37°C. After, samples are then processed again on the gentleMACS Dissociator with two rounds of the “h_impTumor_03.01” program and gently triturated to ensure the tissues are homogenized. Samples are filtered through a 40 μm filter and rinsed with additional sterile-filtered serum-free DMEM. Samples are then centrifuged (1500 rpm, 5 minutes) to pellet the cells and discard the supernatant, then resuspended in ACK lysis buffer and incubated at room temperature for 3-5 minutes. Samples are then diluted with 1X Dubecco’s PBS (1:10), mixed by inverting, and then centrifuged again (1500 rpm, 5 minutes, at 4°C). Finally, cells are resuspended in DPBS, counted, and then centrifuged for a final time before being resuspended in cryopreservation media at 10-20 million live cells per mL. Cells are frozen slowly at -80°C in cryofreezing containers until ready for further downstream processing in a batch with other samples. Samples were prepared for single cell RNA sequencing using standard RNA isolation, cDNA synthesis, and sequencing library preparation protocols according to 10X Genomics protocols and were processed for single-cell sequencing through the Duke Sequencing and Genomic Technologies research core facility.

#### D. Sequencing Analysis

Subsequent single cell RNA sequencing data was analyzed using standard Seurat pipeline to perform QC pre-processing, normalization, identification of highly variable features, scaling, linear dimensionality reduction, and clustering. Cell type identification and annotation was performed using InferCNV analysis (to identify cells with high copy number variations or CNVs, which are likely cancer cells) and SingleR. Uniform manifold approximation and projection (UMAP) plots and heat maps were generated using R studio.

### Clinical Regression Free Survival Analysis

The association between UGDH expression level and recurrence free survival (RFS) in patients with ER positive breast cancer (BC) was assessed via an online Kaplan-Meier plotter (kmplotter.com)^38^. This platform consolidates patient data from The Cancer Genome Atlas (TCGA) project, Gene Expression Omnibus (GEO), and Therapeutic Goods Administration (TGA) and allows for the comparison of RFS for patients with low and high expression of specific proteins and RNA.

### Statistical Analysis

*In vitro* and *in vivo* experiments were primarily analyzed with Student’s t-test, one-way ANOVA, or two-way ANOVA with Bonferroni post-hoc correction. Survival curves were analyzed with the Gehan- Breslow-Wilcoxon test. The threshold of significance was p<0.05 with confidence intervals of 95%.

## AUTHOR CONTRIBUTIONS

- **Meghan J. Price**: substantially contributed to the conception of the study, designed study experiments; performed data acquisition, analysis, and interpretation of results for experiments presented in Figures 1 and 2; aided in interpretation of results for experiments in all figures; drafted the manuscript (original) and provided substantial revisions for its iterations
- **Annee D. Nguyen**: designed experiments; performed data acquisition, analysis, and interpretation of results for experiments presented in Figures 3 and 4; aided in interpretation of results for experiments in all figures; generated figures; drafted portions of the manuscript and provided substantial revisions for the manuscript and its iterations
- **Corinne Haines**: creation of R code used to analyze single-cell RNA-sequencing data from patient samples; performed downstream analysis of sequencing studies (in vitro cell sequencing and human patient single cell sequencing); aided in interpretation of data
- **César D. Baëta**: aided in interpretation of data; consented patient for specimen acquisition and processing for sub-figures in Figure 4
- **Jovita Byemerwa:** aided in data interpretation
- **Debarati Murkajee:** aided in study design and experimental optimization of migration assays; provided minor revisions; aided in interpretation of results for experiments presented in Figures 1 and 2
- **Suzanne Wardell**: contributed to the design and execution of animal studies included in Figures 1-3; aided in interpretation of results of said studies
- **Sandeep Artham**: contributed to the design and execution of animal studies included in Figures 1-3; aided in interpretation of results of said studies
- **Vardhman Kumar**: generated custom microfluidics model, designed & executed *ex vivo* experiment featured in Figure 3A & B
- **Catherine Lavau**: performed data acquisition and analysis of Figures 1F and 2F; aided in interpretation of data; revised manuscript
- **Shyni Varghese**: generated custom microfluidics model featured in Figure 3A & B
- **C. Rory Goodwin**: idea conception for the study, designed study experiments; aided in interpretation of results for all experiments/figures; drafted the manuscript and figures and provided substantial revisions for its iterations; obtained patient specimen and medical history for single-cell sequencing.

---

**Supplementary Figure 1.**
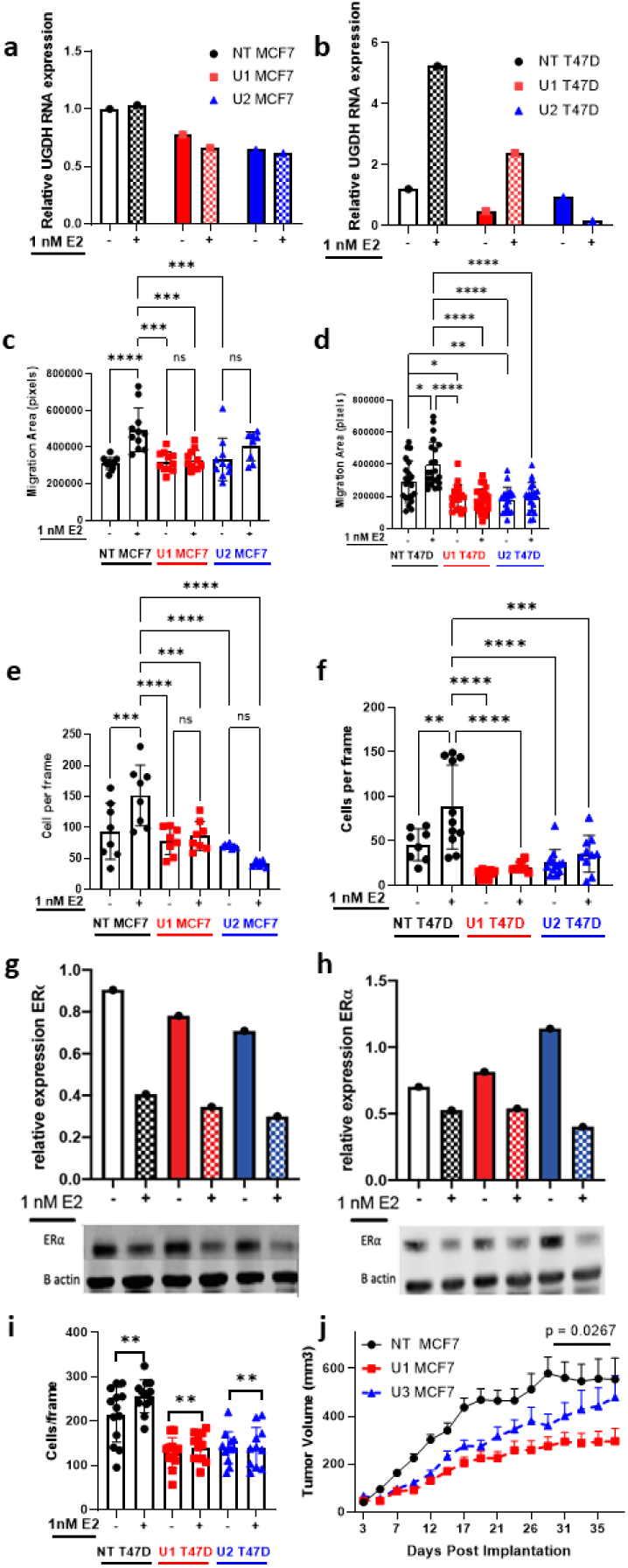
Validation of UGDH shRNA knockdowns and abrogation of migratory phenotypes in ER+ cell lines (MCF7:WS8 and T47D), in two UGDH shRNAs. Black = NT shRNA control cell line, red = UGDH knockdown 1 shRNA, and blue = UGDH knockdown 2 shRNA. **(A)** UGDH RNA expression of MCF7:WS8 and **(B)** T47D treated with (+) or without (-) estrogen (1nM E2). **(C)** Wound healing assays for MCF7:WS8 and **(D)** T47D treated with (+) or without (-) estrogen (1nM E2). . **(E)** Transwell assays for MCF7:WS8 and **(F)** T47D treated with (+) or without (-) estrogen (1nM E2). **(G)** ER-alpha expression of MCF7:WS8 and **(H)** T47D treated with (+) or without (-) estrogen (1nM E2). **(I)** Tumor growth progression of subcutaneous implantation of MCF7:WS8 in mammary fat pad of Nu/J mice implanted with E2 pellets. **(J)** Number of T47D colonies formed on 1% agar after 21 days, with or without E2 stimulation. Statistical analyses were one-way ANOVA for migration assays and colony assays; two-way ANOVA for proliferation assay and tumor growth curve, with Tukey’s post-hoc correction. Data are mean and standard deviation. ns = not significant; *=p<0.05; **=p<0.01; ***=p<0.001; ****=p<0.0001.

**Supplementary Figure 2:**
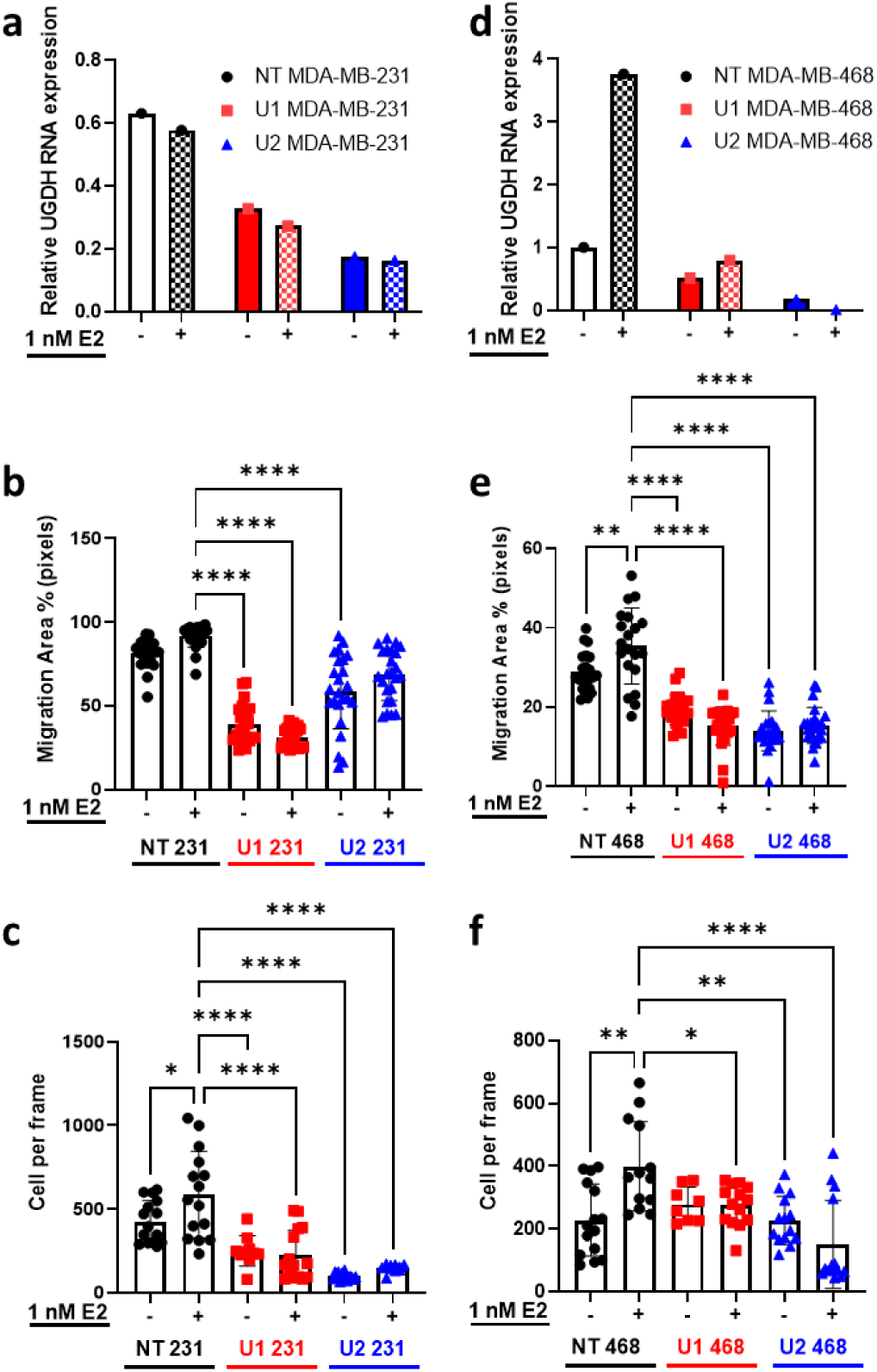
UGDH expression and migration phenotyping in ER negative non-targeting control and shRNA knockdown cell lines, stimulated with and without estrogen. Validation of UGDH shRNA knockdowns and abrogation of migratory phenotypes in ER negative cell lines (MDA-MB-231 and MDA-MB-468), in two UGDH shRNAs. Black = NT shRNA control cell line, red = UGDH knockdown 1 shRNA, and blue = UGDH knockdown 2 shRNA. **(a)** MDA-MB-231 UGDH RNA expression treated with (+) or without (-) estrogen (1nM E2), with **(b)** MDA-MB-231 wound healing assay, and **(c)** MDA-MB-231 transwell assay. **(d)** MDA-MB-468 UGDH RNA expression treated with (+) or without (-) estrogen (1nM E2), with **(e)** MDA-MB-468 wound healing assay, and **(f)** MDA-MB- 468 transwell assay. Data are mean and standard deviation. *=p<0.05; **=p<0.01; ***=p<0.001; ****=p<0.0001.

**Supplementary Figure 3.**
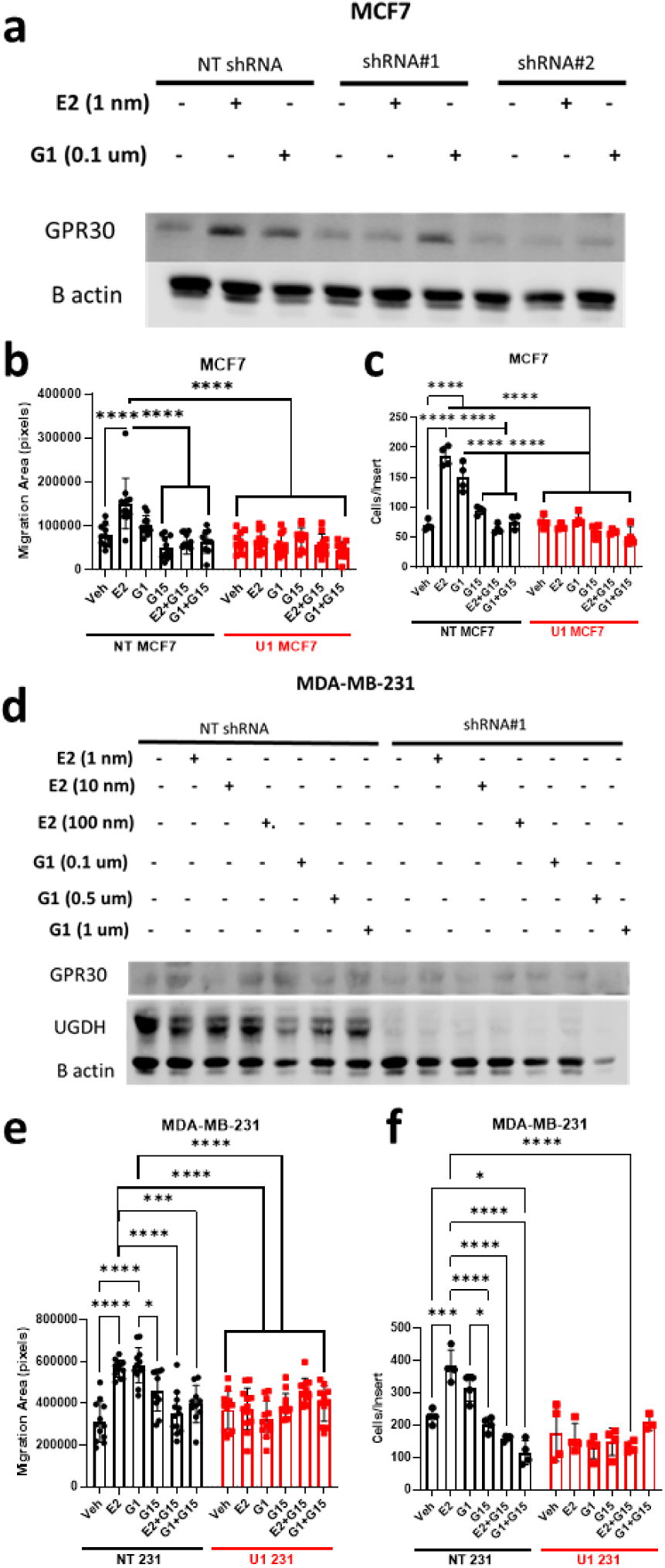
UGDH knockdown prevents GPER agonist (G1)-mediated migration and invasion. Black = NT shRNA control cell line, red = UGDH knockdown 1 shRNA. **(A)** UGDH and GPR30/GPER expression in MCF7:WS8 and **(B)** in MDA-MB-231 stimulated with E2 (1nM, 10 nM, 100 nM) or G1 (0.1uM, 0.5uM, or 1uM). **(C)** Wound healing assay in MCF7:WS8 and **(D)** MDA-MB-231 treated with E2 (1nM), G1 (0.1uM), G15 (GPER antagonist, 1uM), or combination of E2+G15 or G1+G15. **(E)** Transwell invasion assay in MCF7:WS8 and **(F)** MDA-MB-231 treated with E2 (1nM), G1 (0.1uM), G15 (GPER antagonist, 1uM), or combination of E2+G15 or G1+G15. Statistical analyses were one-way ANOVA for migration assays, with Sidak’s post-hoc correction. Data are mean and standard deviation. *=p<0.05; ***=p<0.001; ****=p<0.0001.

**Supplemental Figure 4.**
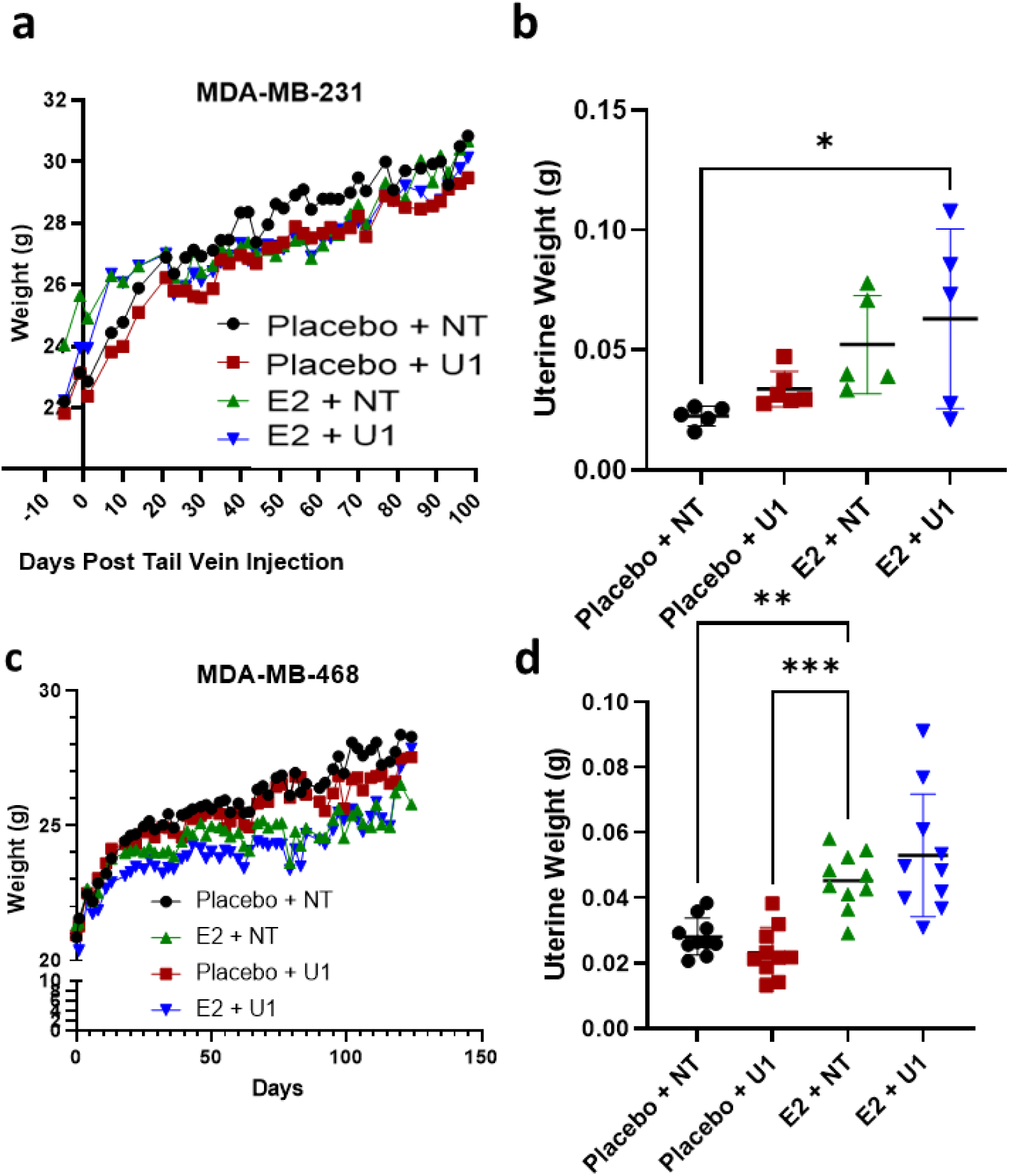
MDA-MB-231 and MDA-MB-468 (ER- cell lines) animal studies. **(A)** Longitudinal weights from surgery date (Day -6) to end of study (Day 100) for MDA-MB-231 metastasis study. **(B)** Uterine weights comparing control cell lines and UGDH KD1 with or without estrogen pellets at end of MDA-MB-231 metastasis study. **(C)** Longitudinal weights for MDA-MB-468 metastasis survival study to end of study (Day 124). **(D)** Uterine weights comparing control cell lines and UGDH KD1 with or without estrogen pellets at end of MDA-MB-468 primary mammary fat pad xenograft study. Statistical analyses were one-way ANOVA with Tukey’s post-hoc correction.

## References

1. Society, A. C. Key Statistics for Breast Cancer, <https://www.cancer.org/cancer/breast-cancer/about/how-common-is-breast-cancer.html> (2022).

2. Karuturi M, V. V., Chavez-MacGregor M. Metastatic breast cancer. In: Kantarjian HM, Wolff RA. The MD anderson manual of medical oncology., Vol. 3e (McGraw-Hill Medical, 2016).

3. Noone AM, H. N., Krapcho M, Miller D, Brest A, Yu M, Ruhl J, Tatalovich Z, Mariotto A, Lewis DR, Chen HS, Feuer EJ, Cronin KA,. SEER Cancer Statistics Review, 1975–2015. (2018).

4 Hsu, J. Y., Chang, C. J. & Cheng, J. S. Survival, treatment regimens and medical costs of women newly diagnosed with metastatic triple-negative breast cancer. Scientific reports 12, 729, doi:10.1038/s41598-021-04316-2 (2022).

5 Gupta, P. B. & Kuperwasser, C. Contributions of estrogen to ER-negative breast tumor growth. J Steroid Biochem Mol Biol 102, 71–78, doi:10.1016/j.jsbmb.2006.09.025 (2006).

6 Gupta, P. B. et al. Systemic stromal effects of estrogen promote the growth of estrogen receptor- negative cancers. Cancer research 67, 2062–2071, doi:10.1158/0008-5472.Can-06-3895 (2007).

7 Sartorius, C. A. et al. Estrogen promotes the brain metastatic colonization of triple negative breast cancer cells via an astrocyte-mediated paracrine mechanism. Oncogene 35, 2881–2892, doi:10.1038/onc.2015.353 (2016).

8 Treeck, O., Schüler-Toprak, S. & Ortmann, O. Estrogen Actions in Triple-Negative Breast Cancer. Cells 9, doi:10.3390/cells9112358 (2020).

9 Friedl, A. & Jordan, V. C. Oestradiol stimulates growth of oestrogen receptor-negative MDA- MB-231 breast cancer cells in immunodeficient mice by reducing cell loss. European journal of cancer **30a**, 1559–1564, doi:10.1016/0959-8049(94)00293-e (1994).

10 Auvinen, P. et al. Hyaluronan in peritumoral stroma and malignant cells associates with breast cancer spreading and predicts survival. The American journal of pathology 156, 529–536, doi:10.1016/s0002-9440(10)64757-8 (2000).

11 Park, J. B., Kwak, H. J. & Lee, S. H. Role of hyaluronan in glioma invasion. Cell Adh Migr 2, 202–207, doi:10.4161/cam.2.3.6320 (2008).

12 Venning, F. A., Wullkopf, L. & Erler, J. T. Targeting ECM Disrupts Cancer Progression. Front Oncol 5, 224, doi:10.3389/fonc.2015.00224 (2015).

13 Zimmer, B. M. et al. Loss of exogenous androgen dependence by prostate tumor cells is associated with elevated glucuronidation potential. Horm Cancer 7, 260–271, doi:10.1007/s12672-016-0268-z (2016).

14 Arnold, J. M. et al. UDP-glucose 6-dehydrogenase regulates hyaluronic acid production and promotes breast cancer progression. Oncogene 39, 3089–3101, doi:10.1038/s41388-019-0885-4 (2020).

15 Oyinlade, O. et al. Targeting UDP-α-D-glucose 6-dehydrogenase inhibits glioblastoma growth and migration. Oncogene 37, 2615–2629, doi:10.1038/s41388-018-0138-y (2018).

16 Lin, L. H. et al. Targeting UDP-glucose dehydrogenase inhibits ovarian cancer growth and metastasis. J Cell Mol Med 24, 11883–11902, doi:10.1111/jcmm.15808 (2020).

17 Wang, X. et al. UDP-glucose accelerates SNAI1 mRNA decay and impairs lung cancer metastasis. Nature 571, 127–131, doi:10.1038/s41586-019-1340-y (2019).

18 Afratis, N. et al. Glycosaminoglycans: key players in cancer cell biology and treatment. Febs j 279, 1177–1197, doi:10.1111/j.1742-4658.2012.08529.x (2012).

19 Lu, P., Takai, K., Weaver, V. M. & Werb, Z. Extracellular matrix degradation and remodeling in development and disease. Cold Spring Harb Perspect Biol 3, doi:10.1101/cshperspect.a005058 (2011).

20 Buijs, J. T. & van der Pluijm, G. Osteotropic cancers: from primary tumor to bone. Cancer Lett 273, 177–193, doi:10.1016/j.canlet.2008.05.044 (2009).

21 Pang, L. et al. Bone Metastasis of Breast Cancer: Molecular Mechanisms and Therapeutic Strategies. Cancers (Basel) 14, doi:10.3390/cancers14235727 (2022).

22 Sun, J. et al. A vertebral skeletal stem cell lineage driving metastasis. Nature 621, 602–609, doi:10.1038/s41586-023-06519-1 (2023).

23 Nguyen, A. D. et al. Single-cell RNA sequencing comparison of the human metastatic prostate spine tumor microenvironment. STAR Protoc 5, 102805, doi:10.1016/j.xpro.2023.102805 (2024).

24 Huang, B., Warner, M. & Gustafsson, J. Estrogen receptors in breast carcinogenesis and endocrine therapy. Mol Cell Endocrinol 418 **Pt** **3**, 240–244, doi:10.1016/j.mce.2014.11.015 (2015).

25 Yaşar, P., Ayaz, G., User, S. D., Güpür, G. & Muyan, M. Molecular mechanism of estrogen- estrogen receptor signaling. Reprod Med Biol 16, 4–20, doi:10.1002/rmb2.12006 (2017).

26 Zhou, K. et al. Estrogen stimulated migration and invasion of estrogen receptor-negative breast cancer cells involves an ezrin-dependent crosstalk between G protein-coupled receptor 30 and estrogen receptor beta signaling. Steroids 111, 113–120, doi:10.1016/j.steroids.2016.01.021 (2016).

27 Masuda, H. et al. Role of epidermal growth factor receptor in breast cancer. Breast Cancer Res Treat 136, 331–345, doi:10.1007/s10549-012-2289-9 (2012).

28 Santen, R. et al. Estrogen mediation of breast tumor formation involves estrogen receptor- dependent, as well as independent, genotoxic effects. Ann N Y Acad Sci 1155, 132–140, doi:10.1111/j.1749-6632.2008.03685.x (2009).

29 Chen, Z. et al. Expressions of ZNF436, β-catenin, EGFR, and CMTM5 in breast cancer and their clinical significances. Eur J Histochem 65, doi:10.4081/ejh.2021.3173 (2021).

30 Kong, X. et al. Mammaglobin, GATA-binding protein 3 (GATA3), and epithelial growth factor receptor (EGFR) expression in different breast cancer subtypes and their clinical significance. Eur J Histochem 66, doi:10.4081/ejh.2022.3315 (2022).

31 Lev, S. Targeted therapy and drug resistance in triple-negative breast cancer: the EGFR axis. Biochem Soc Trans 48, 657–665, doi:10.1042/bst20191055 (2020).

32 Talia, M. et al. The G Protein-Coupled Estrogen Receptor (GPER) Expression Correlates with Pro-Metastatic Pathways in ER-Negative Breast Cancer: A Bioinformatics Analysis. Cells 9, doi:10.3390/cells9030622 (2020).

33. Bin Gao1, P. C., Qifeng Jiang2*. G Protein-Coupled Estrogen Receptor is a Critical Regulator in Metastasis of Breast Cancer Cells. Journal of Biosciences and Medicines, 5, 127–140 (2017).

34 Teoh, S. T., Ogrodzinski, M. P. & Lunt, S. Y. UDP-glucose 6-dehydrogenase knockout impairs migration and decreases in vivo metastatic ability of breast cancer cells. Cancer Lett 492, 21–30, doi:10.1016/j.canlet.2020.07.031 (2020).

35 Wang, T. P., Pan, Y. R., Fu, C. Y. & Chang, H. Y. Down-regulation of UDP-glucose dehydrogenase affects glycosaminoglycans synthesis and motility in HCT-8 colorectal carcinoma cells. Exp Cell Res 316, 2893–2902, doi:10.1016/j.yexcr.2010.07.017 (2010).

36 Giese, A. et al. Dichotomy of astrocytoma migration and proliferation. International journal of cancer 67, 275–282, doi:10.1002/(sici)1097-0215(19960717)67:2<275::Aid-ijc20>3.0.Co;2-9 (1996).

37 Hatzikirou, H., Basanta, D., Simon, M., Schaller, K. & Deutsch, A. ’Go or grow’: the key to the emergence of invasion in tumour progression? Math Med Biol 29, 49–65, doi:10.1093/imammb/dqq011 (2012).

38 Györffy, B. et al. An online survival analysis tool to rapidly assess the effect of 22,277 genes on breast cancer prognosis using microarray data of 1,809 patients. Breast Cancer Res Treat 123, 725–731, doi:10.1007/s10549-009-0674-9 (2010).

38 Zhan, D., Yalcin, F., Ma, D., Fu, Y., Wei, S., Lal, B., Li, Y., Dzaye, O., Laterra, J., Ying, M., Lopez-Bertoni, H., Xia, S. Targeting UDP-a-D-glucose-6 dehydrogenase alters the CNS tumor immune microenvironment and inhibits glioblastoma growth. Genes & Diseases, 9(3): 717–730. doi: 10.1016/j.gendis.2021.08.008. Epub 2021 Sep 17. PMID: 35782977; PMCID: PMC9243400.

39 Chen MB, Whisler JA, Fröse J, Yu C, Shin Y, Kamm RD. On-chip human microvasculature assay for visualization and quantification of tumor cell extravasation dynamics. Nat Protoc. 2017 May;12(5):865–880. doi: 10.1038/nprot.2017.018. Epub 2017 Mar 30. PMID: 28358393; PMCID: PMC5509465.

40 Kumar V, Kinsley D, Perikamana SM, Mogha P, Goodwin CR, Varghese S. Self-assembled innervated vasculature-on-a-chip to study nociception. Biofabrication. 2023 Apr;15(3). doi: 10.1088/1758-5090/acc904.

